# Multimodal non-ionizing quantitative characterization of enamel hypomineralization

**DOI:** 10.64898/2026.01.26.701762

**Authors:** B. Chin, D. Thapa, M. Neshatian, S. Abrams, HG. Moghadam, P.H. Dos Santos, M. Casas, A. Mandelis, L. Bozec

**Affiliations:** Matrix Functionalization and Phenotyping Lab, Faculty of Dentistry, University of Toronto; Centre for Advanced Diffusion-Wave and Photoacoustic Technologies (CADIPT), Department of Mechanical and Industrial Engineering, University of Toronto, Canada; Cliffcrest Dental Office, Scarborough, Canada; Faculty of Dentistry, McGill University, Canada; Department of Paediatric Dentistry, Hospital for Sick Children, Toronto, Canada

**Keywords:** Enamel, Hypomineralization, Molar, Optical Coherence Tomography, Photothermal, Microcomputed tomography, Thermophotonic imaging, lock-in thermography

## Abstract

Enamel hypomineralization is clinically graded but difficult to quantify with visual and radiographic assessment. We evaluated whether optical coherence tomography (OCT) and lock-in thermography imaging (LITI) provide non-ionizing, quantitative lesion localization and phenotyping, using clinical mDDE classification as a practical benchmark and micro-CT (µCT) mineral density as an in vitro reference in a subset. Twenty-five extracted first permanent molars were classified as control, hypomineralized (Type 1 white–cream; Type 2 yellow–brown), or other enamel defects. Co-localized OCT A-scan decay slopes and LITI lock-in phase-contrast were measured in predefined ROIs; six teeth underwent µCT with hydroxyapatite calibration. Clinical scoring showed substantial agreement (inter-rater κ=0.61; intra-rater κ=0.70). OCT decay slopes differed across clinical groups (Kruskal–Wallis p=0.00028), separating controls from defect groups, while separation between hypomineralization and other defects was limited. LITI detected thermophotonic “hot spots” at many clinically identified sites (sensitivity 80%) but showed low specificity (20%) at a single modulation frequency, consistent with high anomaly sensitivity but limited etiologic discrimination. In the µCT subset, hypomineralized ROIs showed reduced mineral density relative to adjacent enamel and spatial concordance with OCT and LITI contrast. Together, OCT+LITI support objective lesion mapping and motivate multi-frequency thermophotonics and expanded reference sampling to improve specificity for chairside translation.

## 1. Introduction

Enamel hypomineralization, commonly referred to as molar hypomineralization (MH)[1] or molar incisor hypomineralization (MIH)[2, 3] is a developmental defect of enamel that arises during amelogenesis. Clinically, it presents as demarcated opacities that range from white-cream to yellow-brown and are frequently accompanied by hypersensitivity, increased caries susceptibility, and post-eruptive breakdown (PEB) [4-6]

Although MH/MIH has historically been associated with perinatal and systemic risk factors [7], current models also support a lesion-centered mechanism in which localized albumin infiltration perturbs enamel maturation during tooth development [8, 9]. With a global prevalence of 13.5% [10], MH/MIH contributes to a substantial clinical burden, including complex restorative needs, treatment challenges in affected teeth, and heightened dental anxiety [5, 11, 12]

In practice, diagnosis and severity grading rely on clinical examination (often supported by radiographs) and indices such as the modified Developmental Defects of Enamel (mDDE) index . While these frameworks standardize descriptive classification, they remain inherently subjective and provide limited quantitative information for stratifying severity or anticipating PEB risk. Conventional radiographs have low sensitivity for hypomineralized enamel changes [13], and their use in children is further constrained by cumulative radiation exposure. A further challenge is that demarcated opacities can overlap visually with early enamel demineralization and other enamel defects, motivating methods that provide objective subsurface contrast. There is therefore a need for objective, reproducible, non-invasive measures that can quantify lesion extent and intensity and that are suitable for longitudinal monitoring. More specifically, quantitative metrics that capture subsurface structure and mineral loss with depth and can be benchmarked against an independent reference standard remain limited for MH/MIH.

Optical and thermo-optical methods are promising candidates for such quantitative phenotyping. Optical coherence tomography (OCT) provides depth-resolved structural contrast and has been widely applied in dentistry, including caries diagnosis[14] and the identification of tissue diseases [15]. OCT has also been explored for developmental enamel defects and hypomineralized enamel, but reproducible quantitative features that are explicitly benchmarked against mineral-density reference measurements remain limited for MH/MIH.

Photothermal radiometry with modulated luminescence (PTR/LUM) has been used primarily for caries detection and monitoring; the Canary System, which incorporates PTR/LUM, can detect non-cavitated caries [16] and recurrent decay [17]. Lock-in thermography imaging (LITI) is the camera-based imaging extension of PTR/LUM and is the thermal modality used in this study. In vitro, microcomputed tomography (µCT) provides high-resolution mineral density mapping that can serve as an in vitro reference for assessing enamel mineral loss, although clinical translation is limited by radiation dose and access.

Here, we assess OCT and LITI for characterizing enamel hypomineralization in extracted first permanent molars. We benchmark imaging findings against expert clinical classification and use µCT to provide spatial and quantitative reference measurements in a subset of specimens. By combining inter- and intra-rater reliability analyses with quantitative signal features derived from OCT and LITI, this study aims to determine whether these modalities can support reproducible, objective assessment of MH/MIH and related enamel defects in a manner consistent with potential clinical translation.

## 2. Materials and Methods

### 2.1 Sample selection, handling, and ethics

Twenty-six extracted first permanent molars (FPMs) were collected from pediatric patients (<18 years) at the University of Toronto Faculty of Dentistry. Teeth were extracted for clinical reasons unrelated to the present study. Informed consent for the collection and research use of extracted teeth was obtained, and ethical approval was granted by the University of Toronto Research Ethics Board (Protocol #00044559). Because enamel hypomineralization requiring extraction is relatively uncommon and ethically cannot be induced, specimen availability is intrinsically limited. The current cohort represents all eligible FPMs collected at the Hospital for Sick Children and the University of Toronto Paediatric Clinic over a 24-month period. As such, the study was designed as an exploratory feasibility investigation rather than a powered diagnostic accuracy trial. Sample size was therefore determined by real-world clinical availability rather than statistical power constraints, consistent with accepted practice for rare-condition imaging studies

Following collection, teeth were disinfected and stored according to protocols adapted from established methods for preserving tissue integrity [13]. Prior to each imaging session, samples were rinsed with distilled water and gently blotted dry to remove surface moisture while minimizing dehydration. Imaging commenced within 30 seconds of blotting; specimens were kept hydrated between modalities to minimise dehydration effects on optical and thermal signals. All specimens underwent imaging in a fixed sequence: standardized photography and radiography, followed by OCT, then LITI. A subset of six molars underwent µCT scanning for mineral density calibration and validation.

### 2.2 Clinical examination

To replicate clinical assessment, standardized photographs were acquired following published guidance for documenting enamel defects using an Olympus OM-D E-M1 camera. Teeth were mounted in dental wax and oriented to obtain consistent views of buccal, lingual/palatal, mesial, and distal surfaces. Radiographs were acquired using size 2 phosphor plates (ScanX, Air Techniques) under standardized settings (70 kV, 125 µA, 0.10 s) to simulate routine diagnostics. Teeth were positioned to approximate a clinical projection geometry. Photographs and radiographs were acquired for documentation and for independent scoring where applicable.

Samples were classified by an expert reference examiner using the modified Developmental Defects of Enamel (mDDE) index (Tab. 1a), which assigns a two-digit code per tooth for defect type and extent (<1/3, 1/3–<2/3, ≥2/3 surface area). mDDE scoring was performed by direct visual examination of the extracted teeth.

Three additional examiners independently scored each tooth using the same mDDE criteria to determine inter-rater reliability (four examiners total). One examiner repeated the assessment after one month to determine intra-rater reliability. Examiners scored independently and were blinded to OCT, LITI, and µCT outputs, and to the other examiners’ ratings.

Based on the reference examiner’s mDDE classification, teeth were assigned to three clinical classification groups: Group 1 (control), no visible enamel defects; Group 2 (hypomineralized), demarcated opacities subcategorized as Type 1 (white–cream) and Type 2 (yellow–brown); and Group 3 (other enamel defects), including diffuse opacities/white spot lesions and nonspecific surface irregularities. Because early caries-related white spot lesions can overlap visually with developmental opacities, this group is treated as a heterogeneous ‘non-MH defect’ comparator rather than a single etiologic category. Clinical classification group labels were finalized a priori from the reference examiner’s mDDE classification before LITI interpretation and before OCT A-scan pattern reliability testing.

### 2.3 Optical coherence tomography (OCT)

For OCT imaging, teeth were blotted dry, mounted on dental wax, and aligned perpendicularly to the incident beam. Buccal, lingual/palatal, mesial, and distal surfaces were scanned from the cementoenamel junction to cusp tip using a VivoSight DX OCT scanner (Michelson Diagnostics, Kent, UK). Occlusal surfaces were excluded due to irregular topography and alignment constraints. Each scan covered a 6 × 6 mm field at 10 kHz, producing 120 B-scans per surface. Individual frames spanned 6000 µm × 1840 µm, with a **lateral sampling of 4.56 µm/pixel**. Images were processed in ImageJ (NIH, USA).

For each non-control tooth, the lesion ROI was defined a priori using the location of the surface-visible defect on standardized photographs. The OCT scan field was positioned to include this defect. The representative B-scan was defined using a rule-based criterion: the B-scan intersecting the midpoint of the lesion’s greatest surface width on that face within the scan window. A-scan profiles were extracted by lateral averaging over a 10-pixel band (≈ 45.6 µm) centered at this location to reduce speckle-related variability. If multiple defects were present on the same surface, the ROI was placed on the defect with the largest surface area (mDDE extent category) and/or the most pronounced discoloration, as recorded during clinical classification. For control teeth, A-scans were extracted from visually sound enamel at an anatomically matched location on the same surface.

A-scans were exported to OriginPro 2025 (OriginLab, USA) for linear regression analysis to quantify signal decay behavior. For each A-scan, intensity was normalized to the maximum at the enamel–air interface peak. The decay segment was defined using an intensity-defined window, bounded by the depths at which the normalized intensity decreased from **90% to 10%** of the surface-peak maximum, constrained to enamel. The enamel extent was determined from the corresponding B-scan (enamel–dentin junction), and the fitting window was truncated at the enamel limit when needed. Linear regression was then performed over this window to compute the decay slope, which was recorded for intergroup comparisons.

### 2.4 Lock-in thermography imaging (LITI)

Lock-in thermography imaging (LITI), the camera-based imaging extension of PTR/LUM, was used as the thermal modality. The system configuration and theoretical background are described elsewhere [18]. Briefly, a square-wave modulated continuous-wave laser (675 nm, 10 W; OsTech Laser, Germany) was used as the excitation source at a 2 Hz modulation frequency. The thermophotonic surface response was recorded using a mid-wave infrared camera (A6700sc, FLIR, USA; 3–5 µm spectral sensitivity) at a 104 Hz sampling rate. The modulation waveform was recorded via a multifunctional data acquisition board.

In-phase (S0) and quadrature (S90) reference signals were generated from the recorded modulation waveform. The thermal sequence was demodulated by weighted averaging to compute pixel-wise amplitude and phase (φ):

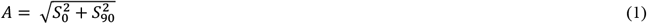

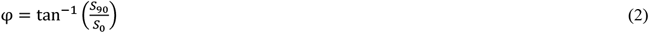

This yielded lock-in amplitude and phase images. For enamel defect analysis and binary anomaly localization, we used a lock-in phase-contrast image (lesion region referenced to surrounding sound enamel within the same field of view) because phase-based contrast is comparatively robust to non-uniform heating and emissivity variation. To extend defect localization toward depth-resolved contrast, Truncated Correlation–Photothermal Coherence Tomography (TC-PCT) was implemented [19], following the dental imaging methodology described in Thapa et al. (2022)[20]. Lesions were classified as LITI-positive when lock-in contrast spatially co-localized with the clinically identified defect region. LITI interpretation was performed blinded to clinical diagnostic group labels. For LITI scoring, the reader was provided only with the tooth identifier and the clinically marked defect location(s) (ROI) derived from the standardized photographs, but not the mDDE-based diagnostic group assignment (control vs hypomineralized vs other defect).

### 2.5 Microcomputed tomography (µCT)

Six FPMs (two per clinical classification group: control, Type 1, and Type 2) were scanned using a NeoScan N80 micro-CT scanner (NeoScan, Belgium). Teeth were blotted dry, mounted in plastic containers with dental wax, and scanned alongside two hydroxyapatite (HA) phantoms (0.238 and 0.749 gHA/cm^3^) for calibration. Scans were performed at 92 kV and 174 µA, with a 0.25 mm copper filter, 7.5 µm voxel size, and 2800 × 2400 detector resolution. A full 360° rotation was acquired with 0.5° steps, drift compensation, and three-frame averaging. Raw projections were reconstructed using NeoScan 80 software. A calibration file derived from HA phantom scans acquired under identical conditions was used to convert grayscale values to mineral density (gHA/cm^3^).

For mineral density quantification, calibrated reconstructions were examined as 2D slices. For each tooth, five slices spanning the lesion region (or the corresponding anatomical region in controls) were selected for analysis. On each slice, enamel mineral density was measured using rectangular (box) regions of interest (ROIs) placed within enamel areas exhibiting distinct grayscale contrast. ROIs were adjusted in size as needed to remain fully within enamel and to avoid cracks/voids, the enamel surface, the dentino-enamel junction, and other boundaries to minimize partial-volume effects. For hypomineralized teeth, ROIs were placed within the clinically identified lesion region, guided by surface photography and the corresponding grayscale contrast on µCT slices; for each slice, a second ROI was placed in adjacent visually sound enamel on the same tooth. Mean mineral density values were computed per ROI for each slice. For hypomineralized teeth, lesion and adjacent-sound ROIs were treated as paired measurements within each slice (five paired slices per tooth). µCT slice selection and ROI placement were not blinded to clinical classification.

### 2.6 Co-registration and ROI definition across modalities (OCT–LITI–µCT)

To enable cross-modality comparison, ROIs were defined on the tooth surface using standardized clinical photographs and then re-identified on OCT and LITI acquisitions using consistent surface landmarks. Teeth were mounted in dental wax in a fixed orientation. Surface landmarks used for alignment included the cementoenamel junction (CEJ), cusp tip(s), line angles (mesial/distal transitions), and, where present, distinctive fissures or restoration margins.

#### OCT ROI definition

OCT scan windows were positioned to encompass the clinically identified defect region. The “lesion-center B-scan” was defined a priori as the B-scan intersecting the midpoint of the lesion’s greatest surface width on that face, as determined from the standardized photograph within the OCT field of view. A-scan profiles were extracted by lateral averaging over a 10-pixel band (≈ 45.6 µm) centered on this location to reduce speckle-related variability.

#### LITI ROI definition and co-localization

LITI was acquired with the same surface facing the excitation/collection axis, and the field of view was positioned using the same surface landmarks to include the clinically identified defect region. Co-localization between OCT and LITI was verified by matching the landmark-referenced defect location (CEJ-to-cusp axis and line angles) across the clinical photograph, the OCT scan window, and the LITI field of view. For Figure 4, OCT and LITI were acquired from the same landmark-matched surface region on each tooth.

#### µCT correspondence (subset)

For µCT specimens, 2D slices were selected to match the lesion region using the same surface anatomy (CEJ-to-cusp direction and lesion position relative to line angles). Lesion and adjacent sound enamel ROIs were placed within enamel, guided by the clinical photograph and µCT grayscale contrast.

### 2.7 Statistical analysis

Statistical analyses were performed using IBM SPSS Statistics for Windows (Version 29.0.1.0, IBM Corp., Armonk, NY, USA). Statistical significance was set at P < 0.05.

- Reliability: Inter-rater agreement for mDDE-based classification (four examiners) and OCT A-scan pattern classification (ten raters) was quantified using Fleiss’ kappa. Intra-rater agreement (repeat scoring by one examiner after one month; and repeat A-scan pattern scoring when applicable) was quantified using Cohen’s kappa.
- OCT regression features: Group differences in OCT A-scan slope were assessed using a Kruskal– Wallis test, followed by Bonferroni-corrected Mann–Whitney U tests for pairwise comparisons.
- µCT mineral density: Paired comparisons of mineral density (lesion vs adjacent sound) were assessed using the Wilcoxon signed-rank test across paired slice-level observations (five paired slices per tooth). Because multiple slices were analyzed per tooth, slice-level comparisons are nested within the specimen and are interpreted as within-tooth evidence in this exploratory subset.
- LITI diagnostic performance: Sensitivity and specificity of LITI were calculated relative to the clinical reference classification, with confidence intervals reported.

Clinical mDDE classification used for diagnostic group assignment was performed by direct visual examination of the extracted teeth. Examiners were blinded to OCT, LITI, and µCT outputs and to other examiners’ ratings. LITI interpretation and OCT A-scan pattern reliability scoring were conducted using image sets blinded to the clinical diagnostic group labels. Quantitative OCT regression features were computed using a fixed processing pipeline with a standardized 90%→10% intensity-defined fitting window. µCT slice selection and ROI placement were performed with reference to the clinically identified lesion region and were not blinded to clinical classification.

This study is exploratory and limited by a modest sample size and a small µCT subset; the heterogeneous ‘other defect’ comparator group; and the single-frequency LITI acquisition, which prioritizes anomaly detection over etiologic specificity. These constraints motivate multi-frequency thermophotonics and expanded reference sampling in future work.

## 3. Results

All samples were photographed and radiographed before advanced imaging analyses. Visual inspection successfully identified enamel opacities consistent with hypomineralization. However, a comparative analysis of clinical photographs and corresponding radiographs (Fig. 1) revealed that radiographic imaging lacked sufficient sensitivity to reliably differentiate hypomineralized from sound enamel. No discernible differences in radiodensity were observed between affected and unaffected regions, highlighting the limitations of conventional dental radiography in detecting qualitative enamel defects.

**Figure 1.**
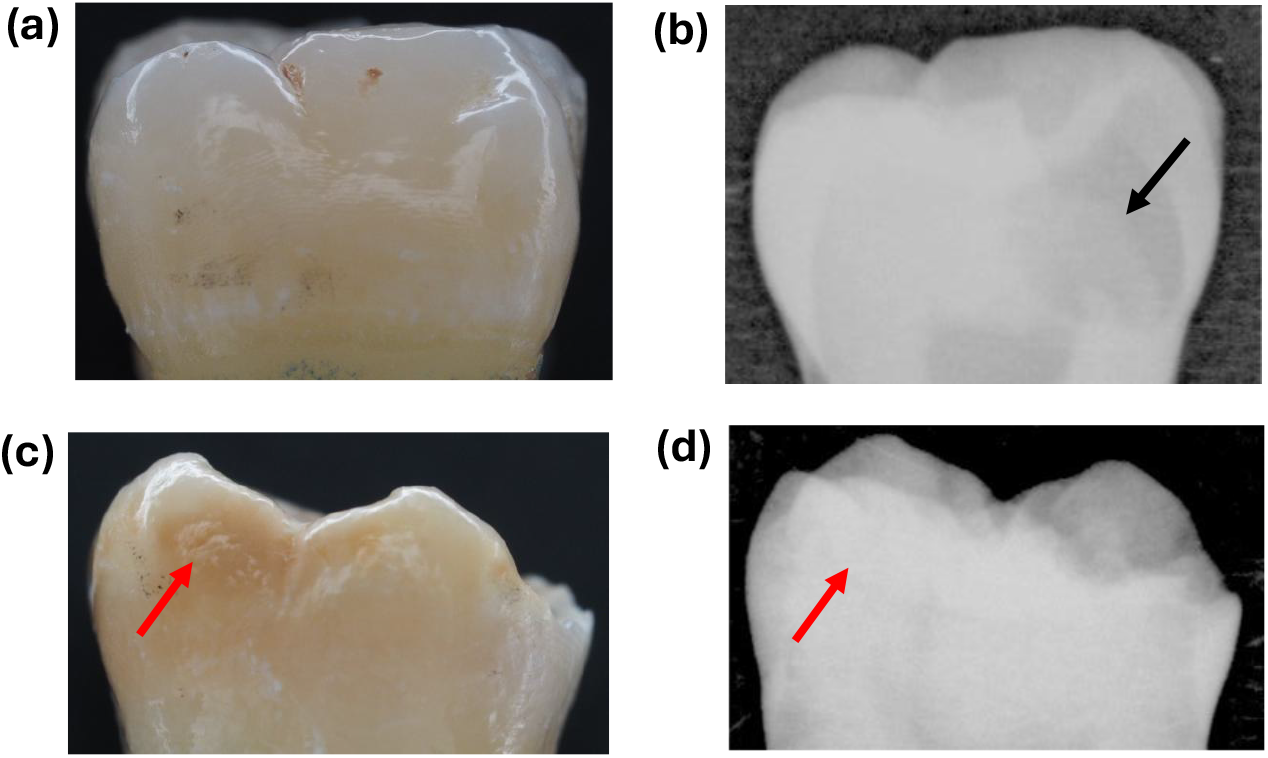
Clinical and radiographic images of a control tooth (a, b), and hypomineralized tooth (c, d). The black arrow in (b) shows a carious lesion not visible clinically, while the red arrow in (c,d) indicates hypomineralization not apparent in (d), underscoring the limited specificity of radiographs for detecting hypomineralization.

### 3.1 Clinical classification

Inter-rater agreement across four examiners was quantified using Fleiss’ κ (κ = 0.61). Intra-rater agreement for repeat scoring by one examiner after one month was quantified using Cohen’s κ (κ = 0.70). Of the 26 FPMs assessed, 25 were included in the final analysis; one specimen was excluded due to enamel loss, which prevented accurate classification. Based on the reference examiner’s mDDE classification (Tab. 1a), teeth were categorized as Group 1 control (n = 7), Group 2 hypomineralized (n = 10; Type 1 n = 6, Type 2 n = 4), and Group 3 other enamel defects (n = 8). The specimen distribution is summarized in Table 1b.

**Table 1.**
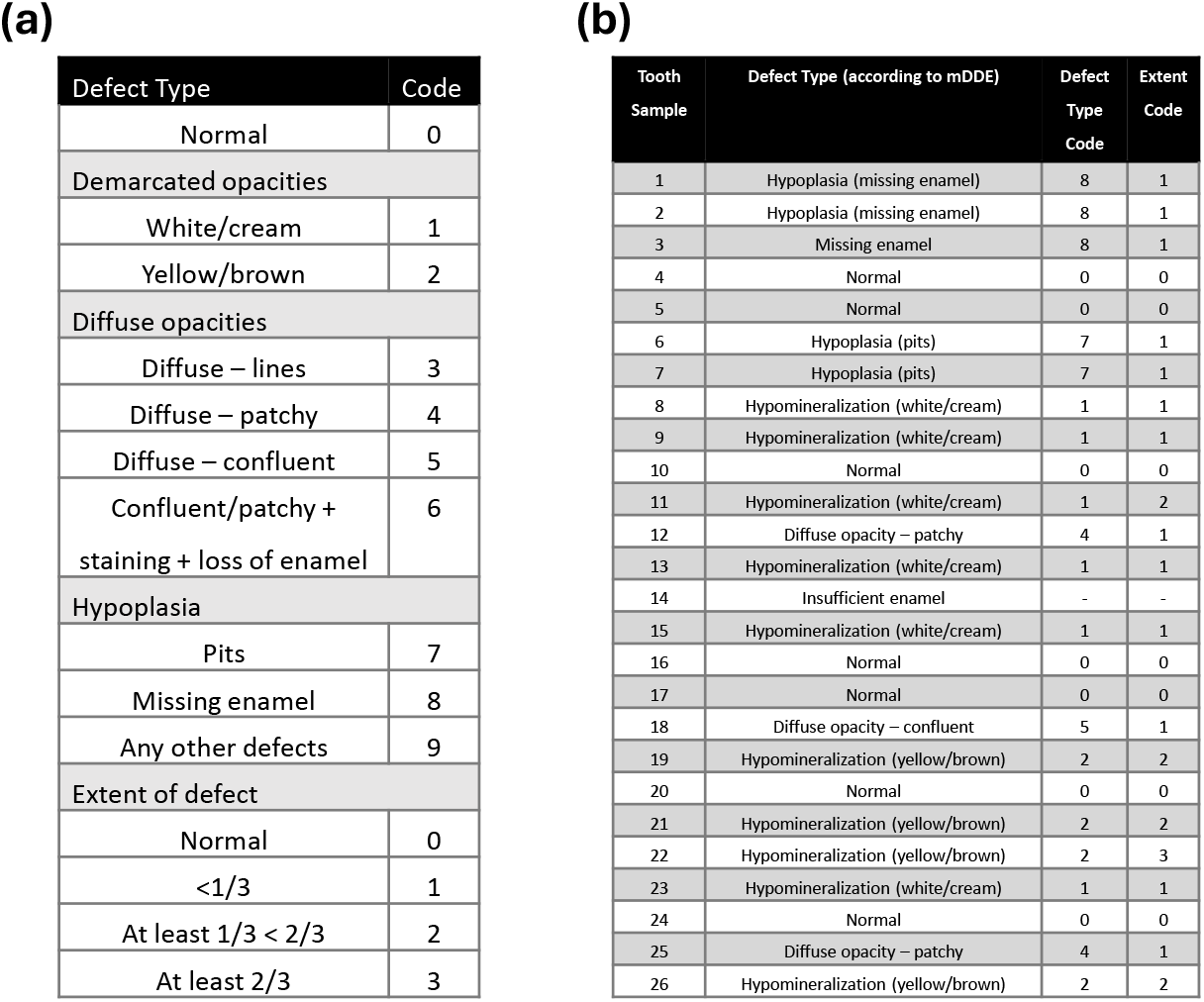

| Defect Type | Code |
| --- | --- |
| Normal | 0 |
| Demarcated opacities |  |
| White/cream | 1 |
| Yellow/brown | 2 |
| Diffuse opacities |  |
| Diffuse – lines | 3 |
| Diffuse – patchy | 4 |
| Diffuse – confluent | 5 |
| Confluent/patchy + staining + loss of enamel | 6 |
| Hypoplasia |  |
| Pits | 7 |
| Missing enamel | 8 |
| Any other defects | 9 |
| Extent of defect |  |
| Normal | 0 |
| <1/3 | 1 |
| At least 1/3 < 2/3 | 2 |
| At least 2/3 | 3 |

(b)
| Tooth Sample | Defect Type (according to mDDE) | Defect Type Code | Extent Code |
| --- | --- | --- | --- |
| 1 | Hypoplasia (missing enamel) | 8 | 1 |
| 2 | Hypoplasia (missing enamel) | 8 | 1 |
| 3 | Missing enamel | 8 | 1 |
| 4 | Normal | 0 | 0 |
| 5 | Normal | 0 | 0 |
| 6 | Hypoplasia (pits) | 7 | 1 |
| 7 | Hypoplasia (pits) | 7 | 1 |
| 8 | Hypomineralization (white/cream) | 1 | 1 |
| 9 | Hypomineralization (white/cream) | 1 | 1 |
| 10 | Normal | 0 | 0 |
| 11 | Hypomineralization (white/cream) | 1 | 2 |
| 12 | Diffuse opacity – patchy | 4 | 1 |
| 13 | Hypomineralization (white/cream) | 1 | 1 |
| 14 | Insufficient enamel | - | - |
| 15 | Hypomineralization (white/cream) | 1 | 1 |
| 16 | Normal | 0 | 0 |
| 17 | Normal | 0 | 0 |
| 18 | Diffuse opacity – confluent | 5 | 1 |
| 19 | Hypomineralization (yellow/brown) | 2 | 2 |
| 20 | Normal | 0 | 0 |
| 21 | Hypomineralization (yellow/brown) | 2 | 2 |
| 22 | Hypomineralization (yellow/brown) | 2 | 3 |
| 23 | Hypomineralization (white/cream) | 1 | 1 |
| 24 | Normal | 0 | 0 |
| 25 | Diffuse opacity – patchy | 4 | 1 |
| 26 | Hypomineralization (yellow/brown) | 2 | 2 |

### 3.2 Qualitative concordance of OCT with µCT

#### 3.2.1 Scan comparison

To evaluate qualitative concordance between OCT and µCT, OCT B-scans were compared with µCT-derived mineral density maps at anatomically matched sites in the µCT subset. Matching was performed qualitatively using the same tooth surface, and landmark-based positioning (CEJ-to-cusp axis and line angles), and the OCT scan position was aligned to the same surface region visualized on reconstructed µCT slices. As shown in Figure 2, lesion regions identified by µCT co-localized with signal anomalies observed in OCT B-scans, showing **qualitative spatial concordance** between OCT structural contrast and regions of reduced mineral density on µCT in the subset examined.

**Figure 2.**
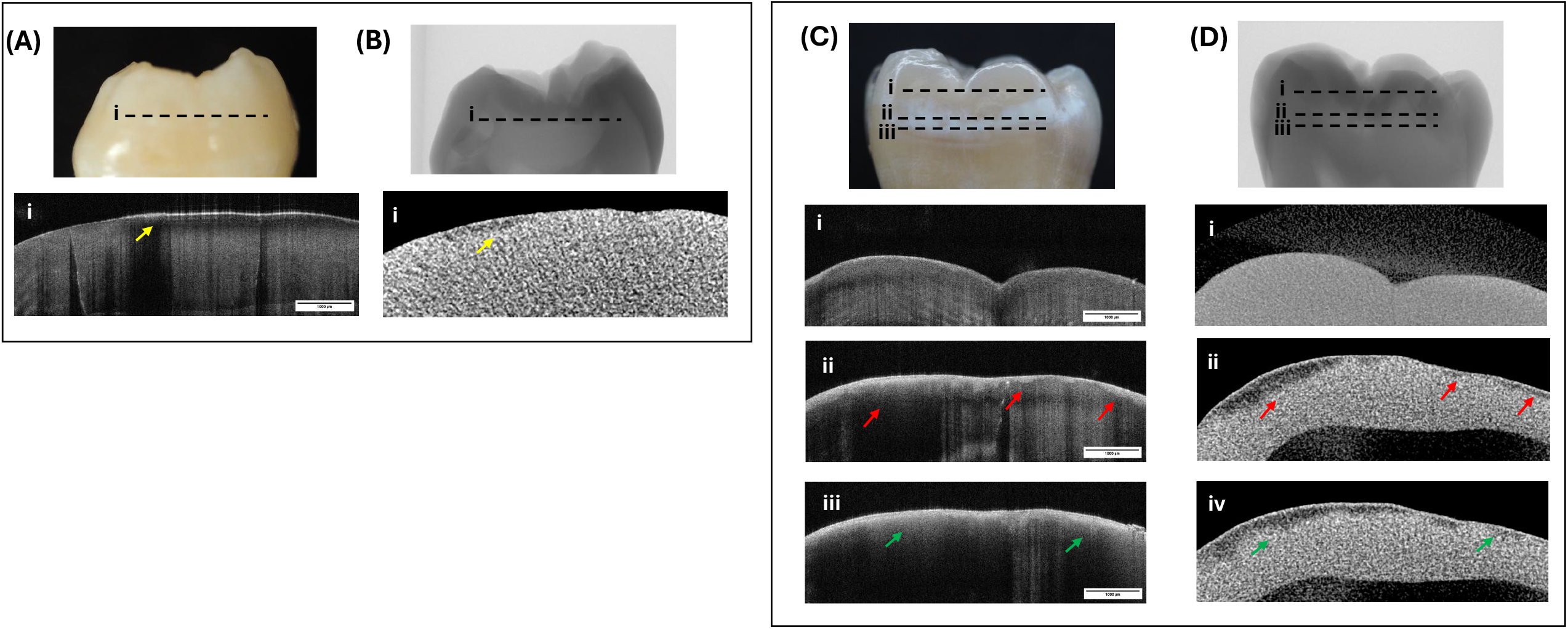
Cross-sectional OCT and µCT imaging of a normal (A) and Type 1 hypomineralized tooth (B). (A) Clinical photo of a sound molar with the OCT scan site marked; the yellow arrow indicates a surface lesion not visible clinically. Corresponding µCT images show a co-localized area of reduced density. i) Clinical image of a molar with a demarcated white-cream opacity. OCT B-scans reveal localized subsurface enamel disruption (red and green arrows), confirmed by µCT slices showing matching areas of reduced mineral density.

#### 3.2.2 Pattern recognition

A-scan signal profiles from representative OCT B-scans revealed five distinct morphologies, characterized by differences in slope, peak count, and peak/base widths. These patterns corresponded to: (1) sound enamel, (2) Type 1 hypomineralization, (3) Type 2 hypomineralization, (4) subsurface enamel disruption not apparent clinically, and (5) diffuse white spot lesions consistent with early caries (Fig. 3). To assess reproducibility, ten calibrated dentists were trained using standardized pattern definitions and classified 35 A-scan images spanning all five patterns; raters were blinded to reference labels. Inter-rater reliability (Fleiss’ κ) was 0.66 and intra-rater reliability was 0.69, indicating substantial agreement.

**Figure 3.**
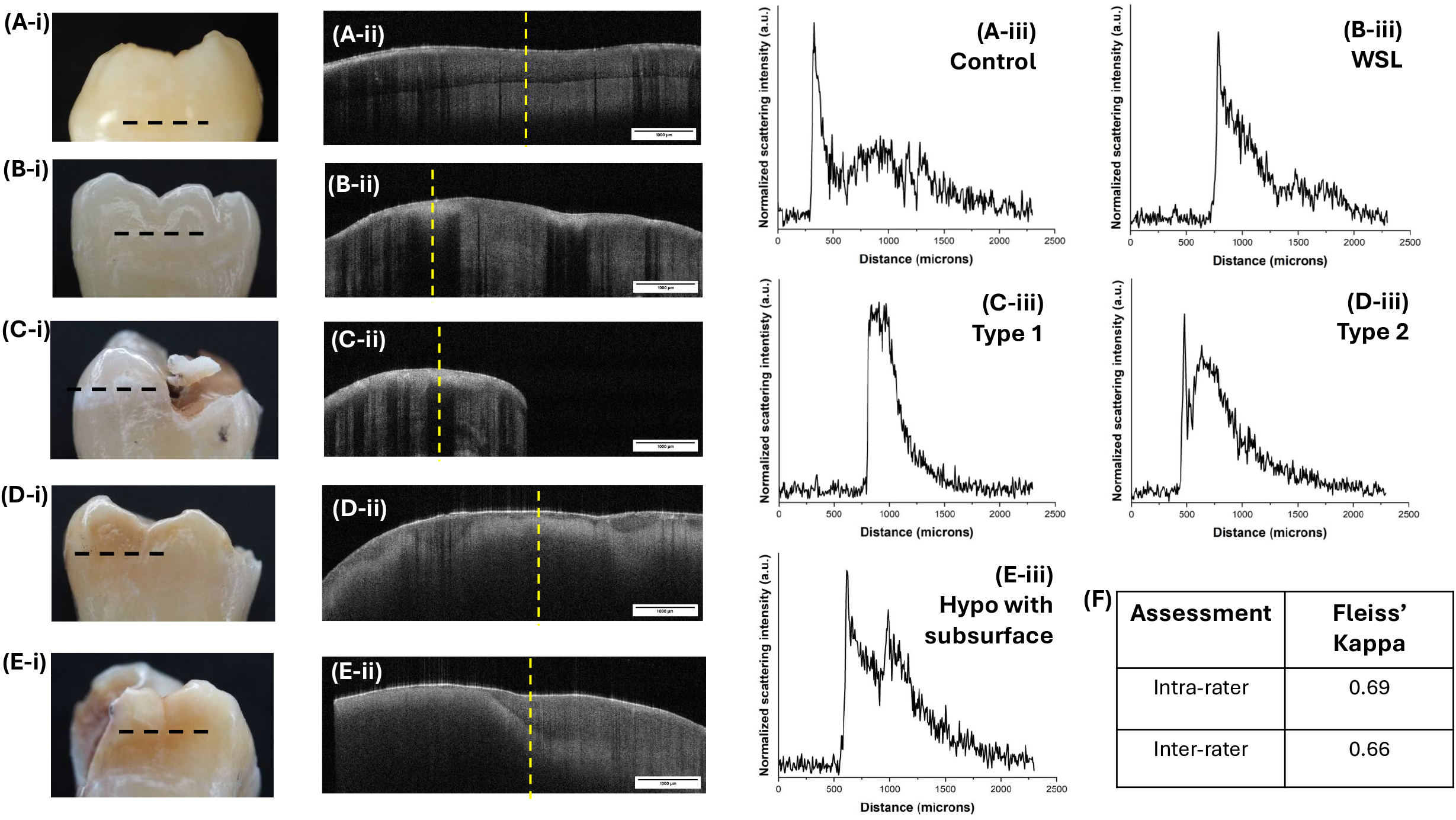
Representative OCT A-scan signal morphologies across enamel conditions. (A–E-i) Clinical images of (A) control enamel, (B) white spot lesion/diffuse opacity, (C) Type 1 (white–cream) hypomineralization, (D) Type 2 (yellow–brown) hypomineralization, and (E) hypomineralization with a subsurface discontinuity. (A–E-ii) Corresponding OCT B-scans; (A–E-iii) A-scans extracted from the indicated regions. Empirically defined A-scan morphologies were characterized by surface peak width/base width, presence of secondary peaks, and post-peak decay shape: (A) sharp surface peak with narrow base and gradual decay; (B) broadened surface peak with narrow base and smooth decline; (C) broadened surface peak with broad base and smooth decay; (D) narrow surface peak with rapid decay and a persistent oscillatory tail; and (E) double-peak profile consistent with a subsurface discontinuity. (F) Fleiss’ kappa showed substantial inter-rater (κ = 0.66) and intra-rater (κ = 0.69) agreement for blinded pattern classification.

### 3.3 Quantitative validation of OCT and µCT

#### 3.3.1 OCT: Linear regression analysis

Linear regression analysis was conducted on 25 A-scan profiles (one per lesion) using OriginPro. Regression windows were defined a priori using a standardized 90%→10% normalized intensity decay segment (within enamel), and slopes were extracted as quantitative descriptors of A-scan signal decay. A Kruskal–Wallis test showed a significant difference in slope values across clinical classification groups (p = 0.00028). Bonferroni-corrected Mann–Whitney U tests confirmed significant differences between control and hypomineralized teeth (p = 0.00031) and between control and other enamel defects (p = 0.0065). The comparison between hypomineralized teeth and other enamel defects did not reach statistical significance after correction (p = 0.062) but showed a trend consistent with gradation in OCT signal behavior across defect categories.

#### 3.3.2 µCT: Mineral density measurement

Mineral density was quantified by µCT in a subset of six teeth (two per diagnostic group). Density values are reported in gHA/cm^3^, derived from hydroxyapatite phantom calibration under identical acquisition and reconstruction conditions. Mean enamel mineral density was 2.87 gHA/cm^3^ for control enamel, 2.23 gHA/cm^3^ for Type 1 hypomineralization, and 2.29 gHA/cm^3^ for Type 2 hypomineralization. Paired slice-level comparisons (lesion vs adjacent sound; five slices per tooth) showed a significant reduction in mineral density at lesion sites relative to adjacent sound enamel (Wilcoxon signed-rank, p = 0.00051). In contrast, comparison of mineral density across clinical classification groups did not show a significant difference (Kruskal–Wallis p = 0.18), consistent with the small sample size and overlapping mineral density profiles across lesion categories.

### 3.4 Lock-in thermography imaging (LITI)

Dynamic thermal-wave imaging was performed at clinically relevant sites using LITI. Lesions were classified as LIT-positive if thermographic signal enhancement corresponded with clinically identified defects. Relative to clinical classification, LITI demonstrated a sensitivity of 80% (95% CI: 49%–94%) and a specificity of 20% (95% CI: 7.0%–45%). At a single modulation frequency (2 Hz), LITI primarily functioned as a sensitive detector of enamel anomalies, motivating multi-frequency acquisition and feature-based classification to improve specificity. To further assess diagnostic performance, LITI and OCT scans were co-localized in two representative teeth exhibiting Type 1 and Type 2 lesions (Fig. 4). While both modalities successfully identified lesion regions, LITI exhibited greater signal intensity in the Type 2 lesion, evidenced by higher contrast and pronounced red coloration in Figure 4B-ii. This suggests enhanced sensitivity of PTR imaging to localized “hot spots” of hypomineralization, potentially reflecting increased organic content or porosity. Among the hypomineralized teeth examined, all four Types 2 lesions exhibited thermographic hot spots, whereas only three of the six Type 1 lesions demonstrated similar findings.

**Figure 4.**
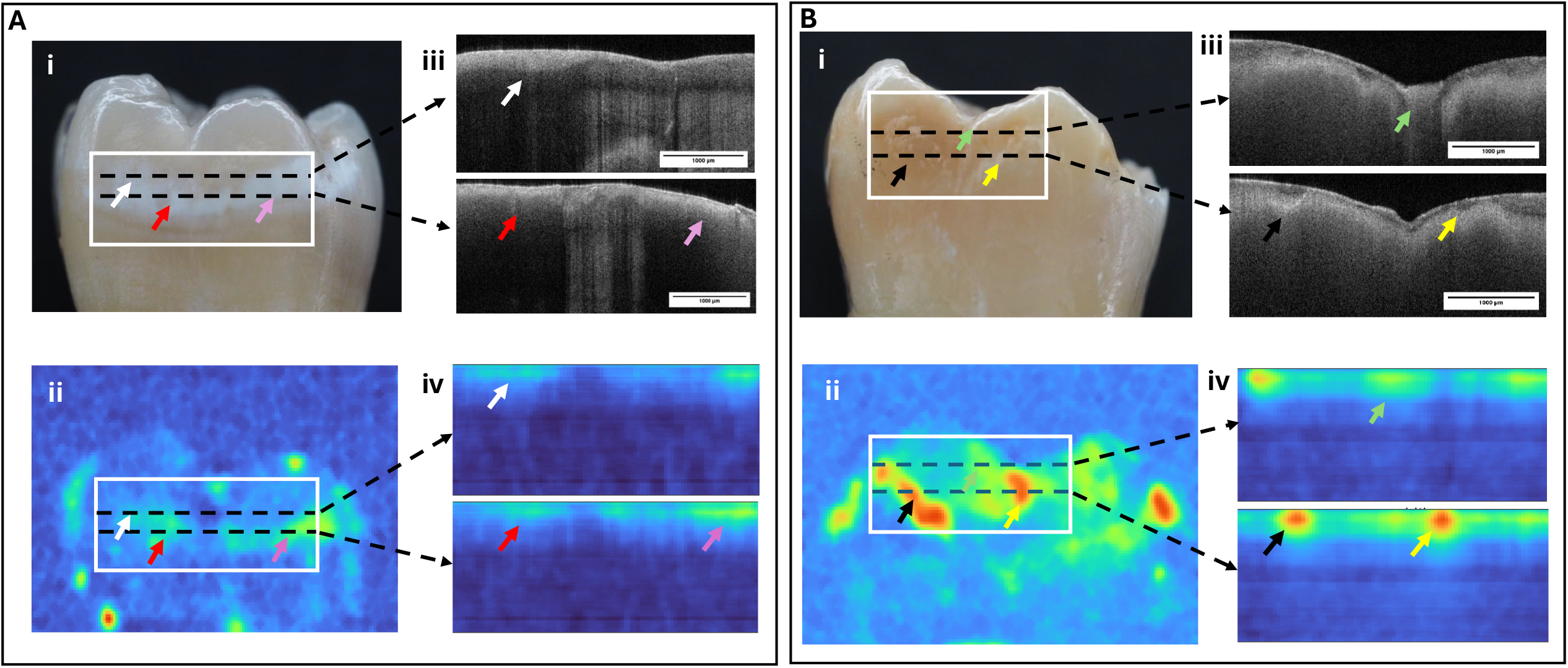
Correlation between LITI and OCT imaging in the evaluation of Type 1 (A) and Type 2 (B) hypomineralization. Fields of view were positioned using CEJ-to-cusp orientation and line-angle landmarks so that the outlined regions corresponded to the same anatomical area across modalities (A-i, B-i) Clinical images of representative lesions. (A-ii, B-ii) LIT surface images (λ=675nm, f=2 Hz) show increased signal in hypomineralized regions. (A-iii, B-iii) Corresponding OCT B-scans from the outlined areas. (A-iv, B-iv) Cross-sectional TC-PCT thermophotonic images reveal lesion depth and extent. Colour-coded arrows highlight co-localized features across modalities. While both lesion types show strong imaging agreement, intense “hot spots” in (B-iv) are not mirrored in (B-iii), suggesting thermal imaging may be sensitive to localized variations not prominent in the corresponding OCT B-scan.

## 4 Discussion

### 4.1 Clinical classification

Enamel hypomineralization is a highly prevalent defect that presents with significant implications. Currently, diagnosis is based on visual or radiographic evaluation [7]. However, as demonstrated in the photographic and radiographic comparison in Figure 1, the findings of this study reinforce the well-established limitation of radiographic diagnosis, namely, its low sensitivity and reliance on subjective interpretation [13]. Radiographic imaging fails to detect subtle mineral and structural changes characteristic of hypomineralization, particularly in its early stages. These limitations underscore the need for objective diagnostic tools that can detect enamel defects before they lead to PEB. The exclusion of one molar from analysis due to extensive enamel loss further underscores the fragility of hypomineralized lesions and the critical importance of early diagnosis.

To complement radiographic assessment, visual classification of all samples was performed using the mDDE index . The index demonstrated substantial inter- and intra-rater agreement, indicating substantial consistency among evaluators. These findings support the utility of the mDDE index in both clinical and epidemiological contexts. While some studies corroborate its reliability [21, 22], others have reported variability in diagnostic accuracy when using this system [23]. Such inconsistencies underscore the limitations of subjective assessment and reinforce the need for objective tools for diagnosing enamel defects.

### 4.2 Tomographic assessment by photon scattering

OCT has been explored as a diagnostic tool in dentistry, particularly for caries [14, 24] and enamel hypomineralization characterization [13, 25-28], making it a key modality for this investigation. In this study, enamel lesions visualized in OCT B-scans closely matched µCT mineral density scans (Fig. 2), supporting OCT as a sensitive modality for detecting enamel defects, with qualitative concordance to µCT in the subset examined. This aligns with findings by de Oliveira et al. (2024)[25] who also reported spatial concordance between OCT-derived contrast patterns and µCT mineral-density findings.

OCT A-scans were used to assess the detection and differentiation of hypomineralization types. A-scans, one-dimensional scattering profiles that map backscattered light intensity as a function of enamel depth [13], provide a direct representation of photon scattering within enamel, where structural defects such as porosity or mineral depletion alter the scattering signal. Building on previously reported empirical A-scan markers [13], we defined five recurring A-scan morphologies based on peak structure and depth-dependent decay (peak width, base width, secondary peaks, and tailing) and used these as a pattern taxonomy to categorize enamel conditions (Fig. 3). Specifically, patterns were operationally defined by: (i) surface peak width and base width, (ii) presence/absence of secondary peaks, and (iii) the shape of the post-peak decay (smooth vs rapid decay with residual tailing). These criteria were documented in a short reference guide and used to train raters before blinded scoring. Because these morphologies are empirical and influenced by acquisition geometry and hydration state, we interpret them as reproducible descriptive signatures rather than unique mechanistic identifiers. Kappa analysis showed substantial inter- and intra-rater agreement, supporting OCT signal patterns as reliable and objective markers for hypomineralization diagnosis. Similar scattering profile patterns for healthy versus hypomineralized teeth were observed by Al-Azri et al. (2016)[13], however, our study is the first to report the reliability of these patterns in lesion classification.

This study identified a novel hypomineralization subtype, “hypomineralization with subsurface lesions”, characterized by a distinct double-peak A-scan pattern (Fig. 3E-iii) suggestive of underlying microfractures. Unlike visual exams, which cannot predict the risk of PEB, this OCT feature may serve as a diagnostic marker of structural vulnerability. Currently, lesion colour, particularly yellow-brown opacities, is the only proposed predictor of severity, though its association with breakdown risk remains unproven [29, 30].

A-scan signal decay trends were used to calculate linear regression slopes as proxies for mineral density, providing an objective characterization of lesion type. Slope values differed significantly between controls and hypomineralized enamel (Bonferroni-corrected Mann–Whitney, p = 0.00031) and between controls and other enamel defects (p = 0.0065), but not between hypomineralization and other defects after correction (p = 0.062). These findings align with prior OCT studies using grayscale values to infer enamel mineral content [31-33]. While methodologies vary, they share the principle that changes in mineralization alter enamel’s optical properties, reflected in OCT signal variation. de Oliveira et al. (2024)[25] evaluated OCT grayscale values to quantify mineral density and reported significant differences in values between healthy and hypomineralized lesions, but did not find differences between mild and severe MH lesions. A-scan slope analysis in this study showed a similar trend, supporting OCT signal analysis as a tool to distinguish mineral density and lesion subtype.

### 4.3 Enamel mineral density by µCT

Mineral density was quantified by µCT in a subset of six teeth (two per diagnostic group) to provide an in vitro reference for mineral density and to support spatial benchmarking of OCT and LITI findings. Mean enamel mineral density was 2.87 gHA/cm^3^ for control enamel, 2.23 gHA/cm^3^ for Type 1 hypomineralization, and 2.29 gHA/cm^3^ for Type 2 hypomineralization. For hypomineralized teeth, mineral density was compared within each tooth by sampling lesion ROIs and adjacent visually sound enamel across five slices spanning the lesion region. Because multiple slices were analyzed per tooth, slice-level measurements are interpreted as within-tooth sampling that captures spatial heterogeneity rather than as independent replicates. Under this tooth-level framing, the data support a consistent within-tooth mineral deficit at lesion sites relative to adjacent enamel (Wilcoxon signed-rank, p = 0.00051), while between-group comparisons across diagnostic categories were not significant in this small subset (Kruskal–Wallis, p = 0.18).

Absolute mineral density values for hypomineralized enamel vary across studies. Neboda et al. (2017)[34] reported mean values of 2.46, 2.21, 1.79, and 2.43 gHA/cm^3^ for unaffected, yellow/creamy, brown, and white enamel, respectively, with significant differences between some lesion categories and substantial overlap for others. Differences between published values and those observed here likely reflect lesion heterogeneity and methodological factors such as calibration, ROI strategy, and within-lesion spatial variability. Collectively, these findings underscore the limitations of generalized lesion categories and support the value of paired within-tooth comparisons and individualized multimodal phenotyping.

### 4.4 Thermophotonic imaging with LITI: sensitivity, specificity, and optimization

LITI, a component of the broader PTR/LUM approach, showed high sensitivity (0.80) for detecting clinically identified defects in this study, but specificity was low (0.20) when acquired at a single modulation frequency. Interpreted at the **tooth/lesion-localization level**, these results indicate that LITI is effective for **anomaly detection and localization**, but is not optimized for discriminating hypomineralization from other enamel defects under the present acquisition setting. This is consistent with thermophotonic contrast reflecting multiple contributors (optical absorption/scattering and thermal transport) that are not unique to MH/MIH. Accordingly, in this single-frequency implementation, LITI is best interpreted as a sensitive screening/localization modality, with a practical route to improving specificity being **multi-frequency acquisition** and feature-based classification leveraging frequency-dependent signatures linked to depth and transport properties. This prospect is consistent with PTR/LUM performance in caries detection, where higher specificity has been reported [35].

In the co-localized examples (Fig. 4), Type 2 lesions exhibited stronger thermographic hot spot contrast than Type 1 lesions, and among hypomineralized teeth examined, all four Type 2 lesions showed hot spots whereas three of six Type 1 lesions did so. Similar trends have been observed in caries studies, where PTR/LUM signal intensity has been associated with lesion severity and increased optical absorption in demineralized regions [36]. Analogous mechanisms may contribute to hypomineralization, where more severe lesions, particularly yellow-brown opacities, may exhibit increased porosity and altered composition, including protein retention [9]. In this context, the thermophotonic hot spot patterns observed here—especially in Type 2 lesions—are consistent with a model in which localized organic-rich regions and/or increased porosity alter optical absorption and enhance non-radiative energy conversion. Because albumin infiltration has been proposed as a lesion-centered mechanism in MH/MIH [8, 9], targeted biochemical validation of high-signal regions (e.g., proteomics) will be required to test whether albumin or other retained proteins contribute to the observed thermophotonic contrast.

While LITI and PTR/LUM are well established in caries detection [16-18, 37, 38], their application to hypomineralization remains comparatively limited. Our findings support the potential of LITI as a sensitive modality for enamel anomaly localization and motivate further optimization for improved specificity in MH/MIH phenotyping.

### 4.5 Combining modalities

To improve diagnostic precision, we adopted a multimodal approach combining LITI with OCT. As shown in Figure 4, regions identified as thermophotonic hot spots in LITI co-localized with OCT structural anomalies on landmark-matched surfaces, supporting tooth-level spatial concordance between enhanced thermophotonic response and subsurface structural alteration. In the µCT subset, this concordance was further supported by within-tooth mineral density reductions at lesion ROIs relative to adjacent enamel, providing an independent reference that strengthens the interpretation that the OCT/LITI contrasts reflect true lesion-associated changes. Importantly, OCT and LITI provide complementary information: OCT offers depth-resolved visualization of lesion boundaries and internal structure based on scattering contrast, whereas LITI highlights thermophotonic heterogeneity that may reflect localized variations in absorption, porosity, and composition.

This co-registered imaging strategy provides a foundation for objective lesion phenotyping that goes beyond purely visual grading. Future work should evaluate multi-frequency thermophotonic acquisition, standardized hydration control, and larger cohorts with expanded µCT reference sampling at the tooth level, with the longer-term aim of translating combined optothermal and tomographic signatures into chairside workflows for early diagnosis and risk-informed clinical management. From a clinical translation perspective, OCT is established as a chairside imaging technology in ophthalmology and has matured into compact, real-time systems in dermatology, illustrating that depth-resolved optical imaging can be deployed in routine workflows. Dentistry presents distinct constraints—limited access, variable surface curvature, specular reflection, hydration dependence, and in vivo motion—yet these are principally engineering and workflow challenges rather than fundamental barriers, and can be addressed through dental-specific handpiece geometries, stabilization approaches, standardized hydration protocols, and real-time quality assurance metrics. In parallel, thermophotonic imaging has a direct translational pathway through existing PTR/LUM platforms; multi-frequency acquisition and quantitative feature extraction are likely to be important for improving specificity. Together, these considerations motivate continued development toward a chairside hybrid system capable of objective lesion phenotyping and longitudinal monitoring in MH/MIH.

## 5 Conclusion

OCT, benchmarked against µCT in a subset of specimens, shows strong potential as a non-invasive approach for objective characterization of enamel hypomineralization. LITI provided complementary contrast by highlighting thermophotonic hot spots that co-localized with defects identified by OCT, supporting its utility for sensitive lesion detection and spatial localization. Together, OCT and LITI offer a multimodal framework that moves beyond purely visual grading toward quantitative lesion phenotyping and longitudinal monitoring.

Future work should focus on improving diagnostic specificity and clinical translation, including multi-frequency thermophotonic acquisition, standardized hydration control, and validation in larger cohorts. Automated analysis approaches, including machine learning for A scan pattern classification and quantitative feature extraction, could support reproducible chairside interpretation. Finally, targeted biochemical validation of high-signal thermophotonic regions, for example, using proteomic assays, may test whether the observed hot-spot contrast is associated with localized protein retention, including albumin, which has been implicated in lesion-centered MH/MIH mechanisms [8, 9]

## 4. Author Contributions

LB conceptualized the work; BC and DT performed data acquisition; BC, DT, MN, HM, PDS, MC, AM, and LB participated in data analysis and interpretation; BC, DT, and LB drafted the manuscript; all authors critically revised the manuscript.

## 5. Data & Code Availability

Data and code are available from the corresponding author upon reasonable request.

## 8. Figure Legends

**1. (a) Modified Developmental Defects of Enamel (mDDE) Index Scoring System**, adapted from Clarkson and O’Mullane (1989), assigning two-digit codes to classify enamel defect type and extent. **(b) Summary of mDDE index scores for all samples**. Of the 26 teeth collected, 25 were analyzed; one was excluded due to extensive enamel loss. Group 1 (n=7) were normal, Group 2 (n=10) hypomineralized (Type 1: n=6; Type 2: n=4), and Group 3 (n=8) showed other defects (e.g., hypoplasia or diffuse opacities). These served as a reference for OCT, LITI, and µCT comparisons.

